## Supplementary Figures for "Convolutional networks can model the functional modulation of the MEG responses associated with feed-forward processes during visual word recognition"

### Supplementary Table 1

Stimuli used in the MEG experiment, the text written inside the stimuli (if any) and the word predicted by the model

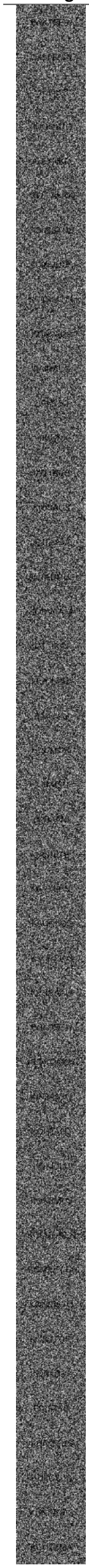

| Stimulus image | Stimulus Type | Stimulus Text | Model prediction |
|---|---|---|---|
| | noisy word | | METSÄTEOLLISUUS |
| | noisy word | | METSÄTEOLLISUUS |
| | noisy word | | METSÄTEOLLISUUS |
| | noisy word | | METSÄTEOLLISUUS |
| | noisy word | | METSÄTEOLLISUUS |
| | noisy word | | METSÄTEOLLISUUS |
| | noisy word | | METSÄTEOLLISUUS |
| | noisy word | | METSÄTEOLLISUUS |
| | noisy word | | METSÄTEOLLISUUS |
| | noisy word | | METSÄTEOLLISUUS |
| | noisy word | | METSÄTEOLLISUUS |
| | noisy word | | METSÄTEOLLISUUS |
| | noisy word | | PÄÄTÖSLAUSELMA |
| | noisy word | | METSÄTEOLLISUUS |
| | noisy word | | METSÄTEOLLISUUS |
| | noisy word | | METSÄTEOLLISUUS |
| | noisy word | | METSÄTEOLLISUUS |
| | noisy word | | METSÄTEOLLISUUS |
| | noisy word | | METSÄTEOLLISUUS |
| | noisy word | | HOMOSEKSUAALISUUS |
| | noisy word | | HOMOSEKSUAALISUUS |
| | noisy word | | METSÄTEOLLISUUS |
| | noisy word | | METSÄTEOLLISUUS |
| | noisy word | | PÄÄTÖSLAUSELMA |
| | noisy word | | METSÄTEOLLISUUS |
| | noisy word | | METSÄTEOLLISUUS |
| | noisy word | | METSÄTEOLLISUUS |
| | noisy word | | METSÄTEOLLISUUS |
| | noisy word | | METSÄTEOLLISUUS |
| | noisy word | | METSÄTEOLLISUUS |
| | noisy word | | PÄÄTÖSLAUSELMA |
| | noisy word | | METSÄTEOLLISUUS |
| | noisy word | | METSÄTEOLLISUUS |
| | noisy word | | METSÄTEOLLISUUS |
| | noisy word | | HOMOSEKSUAALISUUS |
| | noisy word | | OIKEUSKANSLERI |
| | noisy word | | METSÄTEOLLISUUS |
| | noisy word | | METSÄTEOLLISUUS |
| | noisy word | | METSÄTEOLLISUUS |
| | noisy word | | METSÄTEOLLISUUS |
| | noisy word | | METSÄTEOLLISUUS |
| | noisy word | | METSÄTEOLLISUUS |
| | noisy word | | METSÄTEOLLISUUS |
| | noisy word | | METSÄTEOLLISUUS |
| | noisy word | | METSÄTEOLLISUUS |

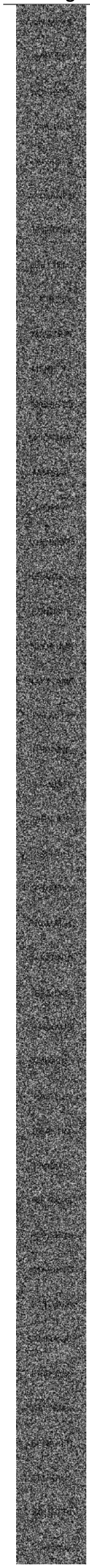

| Stimulus image | Stimulus Type | Stimulus Text | Model prediction |
|---|---|---|---|
| | noisy word | | METSÄTEOLLISUUS |
| | noisy word | | HOMOSEKSUAALISUUS |
| | noisy word | | METSÄTEOLLISUUS |
| | noisy word | | METSÄTEOLLISUUS |
| | noisy word | | METSÄTEOLLISUUS |
| | noisy word | | METSÄTEOLLISUUS |
| | noisy word | | METSÄTEOLLISUUS |
| | noisy word | | METSÄTEOLLISUUS |
| | noisy word | | METSÄTEOLLISUUS |
| | noisy word | | METSÄTEOLLISUUS |
| | noisy word | | METSÄTEOLLISUUS |
| | noisy word | | METSÄTEOLLISUUS |
| | noisy word | | METSÄTEOLLISUUS |
| | noisy word | | METSÄTEOLLISUUS |
| | noisy word | | METSÄTEOLLISUUS |
| | noisy word | | METSÄTEOLLISUUS |
| | noisy word | | METSÄTEOLLISUUS |
| | noisy word | | METSÄTEOLLISUUS |
| | noisy word | | METSÄTEOLLISUUS |
| | noisy word | | METSÄTEOLLISUUS |
| | noisy word | | PÄÄTÖSLAUSELMA |
| | noisy word | | PÄÄTÖSLAUSELMA |
| | noisy word | | METSÄTEOLLISUUS |
| | noisy word | | METSÄTEOLLISUUS |
| | noisy word | | METSÄTEOLLISUUS |
| | noisy word | | METSÄTEOLLISUUS |
| | noisy word | | METSÄTEOLLISUUS |
| | noisy word | | HOMOSEKSUAALISUUS |
| | noisy word | | METSÄTEOLLISUUS |
| | noisy word | | KANSALAISJÄRJESTÖ |
| | noisy word | | TEKNOLOGIATEOLLISUUS |
| | noisy word | | METSÄTEOLLISUUS |
| | noisy word | | METSÄTEOLLISUUS |
| | noisy word | | METSÄTEOLLISUUS |
| | noisy word | | METSÄTEOLLISUUS |
| | noisy word | | METSÄTEOLLISUUS |
| | noisy word | | METSÄTEOLLISUUS |
| | noisy word | | METSÄTEOLLISUUS |
| | noisy word | | HOMOSEKSUAALISUUS |
| | noisy word | | METSÄTEOLLISUUS |
| | noisy word | | HOMOSEKSUAALISUUS |
| | noisy word | | METSÄTEOLLISUUS |
| | noisy word | | METSÄTEOLLISUUS |
| | noisy word | | METSÄTEOLLISUUS |
| | noisy word | | METSÄTEOLLISUUS |

| Stimulus image | Stimulus Type | Stimulus Text | Model prediction |
|---|---|---|---|
| | noisy word | | METSÄTEOLLISUUS |
| | noisy word | | METSÄTEOLLISUUS |
| | noisy word | | METSÄTEOLLISUUS |
| | noisy word | | METSÄTEOLLISUUS |
| | noisy word | | METSÄTEOLLISUUS |
| | noisy word | | METSÄTEOLLISUUS |
| | noisy word | | METSÄTEOLLISUUS |
| | noisy word | | METSÄTEOLLISUUS |
| | noisy word | | METSÄTEOLLISUUS |
| | noisy word | | METSÄTEOLLISUUS |
| | noisy word | | METSÄTEOLLISUUS |
| | noisy word | | METSÄTEOLLISUUS |
| | noisy word | | METSÄTEOLLISUUS |
| | noisy word | | METSÄTEOLLISUUS |
| | noisy word | | METSÄTEOLLISUUS |
| | noisy word | | METSÄTEOLLISUUS |
| | noisy word | | METSÄTEOLLISUUS |
| | noisy word | | METSÄTEOLLISUUS |
| | noisy word | | HOMOSEKSUAALISUUS |
| | noisy word | | METSÄTEOLLISUUS |
| | noisy word | | METSÄTEOLLISUUS |
| | noisy word | | METSÄTEOLLISUUS |
| | noisy word | | METSÄTEOLLISUUS |
| | noisy word | | PÄÄTÖSLAUSELMA |
| | noisy word | | METSÄTEOLLISUUS |
| | noisy word | | HOMOSEKSUAALISUUS |
| | noisy word | | METSÄTEOLLISUUS |
| SSKSRLRT | consonants | SSKSRLRT | LOOGISESTI |
| NKTMPVTN | consonants | NKTMPVTN | KUNTAYHTYMÄ |
| RKSJMTT | consonants | RKSJMTT | ENGLANTI |
| KTSMTRKK | consonants | KTSMTRKK | RETORIIKKA |
| HTRJKSLK | consonants | HTRJKSLK | HYVÄKSYÄ |
| SNKTLTJV | consonants | SNKTLTJV | JONOTTAA |
| HHTNNTSR | consonants | HHTNNTSR | TÄYTÄNTÖÖN |
| TTLKKNHD | consonants | TTLKKNHD | TUTKIMUS |
| TMVHJSPM | consonants | TMVHJSPM | UNIVERSUMI |
| SKJLTRST | consonants | SKJLTRST | VIRALLISESTI |
| RPTTMTHV | consonants | RPTTMTHV | HEITTÄYTYÄ |
| JSSPVMTK | consonants | JSSPVMTK | OPETUSSUUNNITELMA |
| LMTMJTNS | consonants | LMTMJTNS | LAIMINLYÖNTI |
| SSKRNTKR | consonants | SSKRNTKR | SEREMONIA |
| MNSLSRN | consonants | MNSLSRN | AMATÖÖRI |
| KSMNRTT | consonants | KSMNRTT | REMONTTI |
| SVTLNRT | consonants | SVTLNRT | PETTYMYS |
| PMSMHVS | consonants | PMSMHVS | VÄHENNYS |
| STMJVHV | consonants | STMJVHV | TYNNYRI |
| TTNRNHR | consonants | TTNRNHR | UUTINENKIRJE |
| SKKNTMT | consonants | SKKNTMT | SUKUNIMI |
| RLTHLTTR | consonants | RLTHLTTR | VETÄYTYÄ |
| KJLJTRKR | consonants | KJLJTRKR | ÄLYLLINEN |
| SNNTNMRK | consonants | SNNTNMRK | ENNEMMIN |
| VNSHKHKV | consonants | VNSHKHKV | VAROVAINEN |
| RNTKKRVL | consonants | RNTKKRVL | NYKYHETKI |
| NKMNMLTH | consonants | NKMNMLTH | MUMMU |
| SSPRRPKT | consonants | SSPRRPKT | SERVERI |
| MMTTVHSL | consonants | MMTTVHSL | MUTTAEI |
| NLRNVNSR | consonants | NLRNVNSR | VALMENNUS |
| VRSTTMTH | consonants | VRSTTMTH | OPPITUNTI |
| SLPSRKSR | consonants | SLPSRKSR | YLEISRADIO |
| SPLTSKGS | consonants | SPLTSKGS | PEITOSSA |
| HRTKNNTR | consonants | HRTKNNTR | TÄYDENNYS |
| SRLMPSNL | consonants | SRLMPSNL | ERIVÄRINEN |
| VVHDTKSS | consonants | VVHDTKSS | HYPOTEESI |
| PMRSRNVK | consonants | PMRSRNVK | JOKSEENKIN |
| KHHTVKNJ | consonants | KHHTVKNJ | RUHTINAS |
| SRKSRVKK | consonants | SRKSRVKK | JOKSEENKIN |
| KVKTRHTV | consonants | KVKTRHTV | MOOTTORITIE |
| MHMMVTTN | consonants | MHMMVTTN | VAKIINNUTTAA |
| TTPTNSTT | consonants | TTPTNSTT | LIITTOVALTIO |
| SMLVKMKR | consonants | SMLVKMKR | HAKULOMAKE |
| TMLMTJKR | consonants | TMLMTJKR | OMINAINEN |
| RTKHLSTT | consonants | RTKHLSTT | VIOLETTI |
| MKJRKSHV | consonants | MKJRKSHV | NÄKÖKOHTA |
| RKJTTSNN | consonants | RKJTTSNN | IHOTTUMA |
| LRRTHPSJ | consonants | LRRTHPSJ | LÄHEISESTI |
| RSTTSSTS | consonants | RSTTSSTS | REILUSTI |
| MMTHKNNT | consonants | MMTHKNNT | OMITUINEN |
| NHNTVTSM | consonants | NHNTVTSM | NIMITTÄIN |
| RVGTSKPT | consonants | RVGTSKPT | PROJEKTI |
| TLRTTKMT | consonants | TLRTTKMT | TYÖTUNTI |
| SHHPNNTT | consonants | SHHPNNTT | HORISONTTI |
| LSSKKNKH | consonants | LSSKKNKH | ESIKOINEN |
| LNLRSSNN | consonants | LNLRSSNN | DIVISIOONA |
| KMKLTSKL | consonants | KMKLTSKL | VÄKILUKU |
| KTTNKNMK | consonants | KTTNKNMK | HALLINNONALA |
| NVLSNSVL | consonants | NVLSNSVL | MYRKKY |
| STGLTHDJ | consonants | STGLTHDJ | PELOTTAVA |
| RSMMPTRL | consonants | RSMMPTRL | EPÄONNISTUA |
| RSHLTVKM | consonants | RSHLTVKM | EHDOLLINEN |
| JNPTKLST | consonants | JNPTKLST | METALLI |
| FVRLSTKR | consonants | FVRLSTKR | TRAILERI |
| KJSMRTK | consonants | KJSMRTK | KUUMETA |
| NLJRVSK | consonants | NLJRVSK | KATKERA |
| JMNKTSP | consonants | JMNKTSP | VANKEUS |
| SLNPRTS | consonants | SLNPRTS | PUMPPU |
| THLSRVN | consonants | THLSRVN | TULOSTIN |
| THTRMKT | consonants | THTRMKT | TYÖTUNTI |
| VVNJTSR | consonants | VVNJTSR | WHICH |
| JRVSNKH | consonants | JRVSNKH | TÄYDENNYS |
| PKNSHMN | consonants | PKNSHMN | LEMPINIMI |
| MHVDDMST | consonants | MHVDDMST | MUUSIKKO |
| NLTSVNL | consonants | NLTSVNL | VÄLILYÖNTI |

| Stimulus image | Stimulus Type | Stimulus Text | Model prediction |
|---|---|---|---|
| SKNJTPT | consonants | SKNJTPT | SANOTTU |
| HSSVTST | consonants | HSSVTST | RESEPTI |
| TPPLJKK | consonants | TPPLJKK | TEHTÄVÄ |
| LLKJJLM | consonants | LLKJILM | YLÄKULMA |
| PSVNRTV | consonants | PSVNRTV | SYVENTYÄ |
| TPTKHPN | consonants | TPTKHPN | TYÖVAIHE |
| SRNPTRS | consonants | SRNPTRS | ROKOTUS |
| PTRHRJT | consonants | PTRHRJT | ETUPUOLI |
| TBMRLNS | consonants | TBMRLNS | TEMPAUS |
| VSMNKTSH | consonants | VSMNKTSH | ENEMMISTÖ |
| NRLTSTK | consonants | NRLTSTK | MIELTYÄ |
| JTKRVRT | consonants | JTKRVRT | TUOPPI |
| LSLLRVS | consonants | LSILRVS | LISTAUS |
| RTSMSKK | consonants | RTSMSKK | RETORIIKKA |
| LLRSSLS | consonants | [LRSSIS | CLASSIC |
| KNTKRPR | consonants | KNTKRPR | MELKEEN |
| SBTRTTN | consonants | SBTRTTN | ESTEETÖN |
| TNTLVSK | consonants | TNTLVSK | TUNTUINEN |
| MSSMSRN | consonants | MSSMSRN | MIGREENI |
| HNHKSTR | consonants | HNHKSTR | YMPÄRISTÖ |
| MTSMTSTR | consonants | MTSMTSTR | ÄLYKKYYS |
| NLTSRKT | consonants | NLTSRKT | INTERNET |
| TNSRVNV | consonants | TNSRVNV | LEMPINIMI |
| JTPLKRN | consonants | JTPLKRN | TIETÄEN |
| LSLHLNT | consonants | LSLHLNT | PELISÄÄNTÖ |
| HMLKLNN | consonants | HMLKLNN | JUMALAINEN |
| VSHMMTK | consonants | VSHMMTK | LOPPUUNMYYDÄ |
| HTVVPRS | consonants | HTVVPRS | VYÖHYKE |
| NTRNTLS | consonants | NTRNTLS | KYYNELE |
| HLTRNNM | consonants | HLTRNNM | KUITENKIN |
| KTTKVHS | consonants | KTTKVHS | VIITEKEHYS |
| LHTPRMTT | consonants | LHTPRMTT | YHTEYDENOTTO |
| LRTSNRN | consonants | LRTSNRN | TIETENKIN |
| HLPPTMK | consonants | HLPPTMK | RIIPPUMA |
| JTTKNRM | consonants | JTTKNRM | TYTTÖNEN |
| RNLKLRK | consonants | RNLKLRK | PIKALAINA |
| KTRNSLR | consonants | KTRNSLR | FLUNSSA |
| JRDNSST | consonants | JRDNSST | PRONSSI |
| RKTPNNB | consonants | RKTPNNB | ANTENNI |
| DMHMSHR | consonants | DMHMSHR | DOMINOIDA |
| TNKRLST | consonants | TNKRLST | FINALISTI |
| MSRRKKS | consonants | MSRRKKS | NUOREKAS |
| AHMAAJA | pseudoword | AHMAAJA | AINAKAAN |
| HAIKULI | pseudoword | HAIKULI | JÄÄHALLI |
| ÖYTÖNTÖ | pseudoword | ÖYTÖNTÖ | ISTUNTO |
| SAALESTO | pseudoword | SAALESTO | SÄÄTELY |
| KAALETAS | pseudoword | KAALETAS | MAALAUS |
| SEPIVUUS | pseudoword | SEPIVUUS | SOPIVUUS |
| TORJONTA | pseudoword | TORJONTA | TORJUNTA |
| TANNAALI | pseudoword | TANNAALI | TANKATA |
| SAHTOORI | pseudoword | SAHTOORI | BAKTEERI |
| KAARESTO | pseudoword | KAARESTO | KAHDESTI |
| TANTEERI | pseudoword | TANTEERI | PANTTERI |
| VIKKEEJA | pseudoword | VIKKEEJA | ILMISELVÄ |
| HANKANTA | pseudoword | HANKANTA | UIMARANTA |
| VOOSTAUS | pseudoword | VOOSTAUS | VUOSITASO |
| KELIISSI | pseudoword | KELIISSI | KULISSI |
| KAASENTA | pseudoword | KAASENTA | KARSINTA |
| SUUNISTO | pseudoword | SUUNISTO | TUTKIMUSTYÖ |
| KAUSAATA | pseudoword | KAUSAATA | KOODATA |
| KIISSETTA | pseudoword | KIISSETTÄ | YLEISTYÄ |
| RIHAAMI | pseudoword | RIHAAMI | OHJAAMO |
| JUUMARI | pseudoword | JUUMARI | DUUNARI |
| KIKOUMA | pseudoword | KIKOUMA | KUOLEMA |
| HEIKÄLE | pseudoword | HEIKÄLE | YHTÄÄLTÄ |
| HERVUNTA | pseudoword | HERVUNTA | HYÖDYNTÄÄ |
| HESPELI | pseudoword | HESPELI | UPEASTI |
| HIEVERÄ | pseudoword | HIEVERÄ | RIIPPUA |
| HILLIMÄ | pseudoword | HILLIMÄ | VÄITTÄMÄ |
| HOPORKA | pseudoword | HOPORKA | KIRSIKKA |
| HOVATTA | pseudoword | HOVATTA | PALVELLA |
| LIHMAAJA | pseudoword | LIHMAAJA | TYÖMÄÄRÄ |
| AISTAITA | pseudoword | AISTAITA | SATSATA |
| INKRIHTI | pseudoword | INKRIHTI | EMOLEVY |
| JAHNAAJA | pseudoword | JAHNAAJA | SAARNAAJA |
| JAAKOUTA | pseudoword | JAAKOUTA | LAUKAISTA |
| JARNUUMO | pseudoword | JARNUUMO | SÄHKÖINEN |
| JÄMÄHJE | pseudoword | JÄMÄHJE | LIMAKALVO |
| JÖRSITE | pseudoword | JÖRSITE | PAKETTI |
| KEHKÄNTÄ | pseudoword | KEHKÄNTÄ | RUOKAHALU |
| KEIJAKE | pseudoword | KEIJAKE | KOLIKKO |
| KEENETSI | pseudoword | KEENETSI | KESKELLE |
| KEPERNO | pseudoword | KEPERNO | KYSEINEN |
| AMPAUTA | pseudoword | AMPAUTA | VÄKIVALTA |
| KIHELAS | pseudoword | KIHELAS | KIVULIAS |
| LEMMOSTI | pseudoword | LEMMOSTI | TUNNISTE |
| LEUHAKE | pseudoword | LEUHAKE | TELKKARI |
| LIEHENNE | pseudoword | LIEHENNE | TIETOINEN |
| LIEKÄNÄ | pseudoword | LIEKÄNÄ | LUPAAVA |
| LIHTAUTA | pseudoword | LIHTAUTA | TALVISOTA |
| LIPALKO | pseudoword | LIPALKO | TILIKAUSI |
| LUMUTTI | pseudoword | LUMUTTI | LÄHETTI |
| MAUHAATA | pseudoword | MAUHAATA | MATKAILU |
| MAUKUTU | pseudoword | MAUKUTU | MÄÄRÄYTYÄ |
| ANSAUTA | pseudoword | ANSAUTA | ANSAITA |
| MEJAAKA | pseudoword | MEJAAKA | MAJAKKA |
| MIEHATA | pseudoword | MIEHATA | MUURATA |
| MYÖHÄTÄ | pseudoword | MYÖHÄTÄ | MYÖTÄILLÄ |
| MÖLSÄTÄ | pseudoword | MÖLSÄTÄ | MELODIA |
| NAKLAATA | pseudoword | NAKLAATA | MAKUASIA |
| NENANTI | pseudoword | NENANTI | REMONTTI |
| NIHKAUTA | pseudoword | NIHKAUTA | MYÖHÄSTYÄ |

| Stimulus image | Stimulus Type | Stimulus Text | Model prediction |
|---|---|---|---|
| NILKOVA | pseudoword | NILKOVA | MITÄTÖN |
| NORJAMA | pseudoword | NORJAMA | KORJAAMO |
| OHJISTU | pseudoword | OHJISTU | TAISTELU |
| AAPRASKA | pseudoword | AAPRASKA | HALKEAMA |
| OOHTAISI | pseudoword | OOHTAISI | LEHTITALO |
| OISTAARI | pseudoword | OISTAARI | FESTARI |
| PAAKKELE | pseudoword | PAAKKELE | IHANASTI |
| PAHAKSU | pseudoword | PAHAKSU | PÄIVÄKOTI |
| PAATALJA | pseudoword | PAATALJA | OHJAAJA |
| PEHNAAJA | pseudoword | PEHNAAJA | SEURAAJA |
| PEIJAMA | pseudoword | PEIJAMA | PELIAIKA |
| PESLAAKI | pseudoword | PESLAAKI | YÖELÄMÄ |
| PIIMIKE | pseudoword | PIIMIKE | SILMÄYS |
| PIISASMA | pseudoword | PIISASMA | YLÖSPÄIN |
| ELNUKKA | pseudoword | ELNUKKA | SILMUKKA |
| PINVIITA | pseudoword | PINVIITA | HYMYILLÄ |
| POHVAAJA | pseudoword | POHVAAJA | EPÄVAKAA |
| POKOSTA | pseudoword | POKOSTA | PURSUTA |
| PÄHLÄKE | pseudoword | PÄHLÄKE | PÄÄLAKI |
| PÖHKERÖ | pseudoword | PÖHKERÖ | POIKKEUS |
| RAAHISTE | pseudoword | RAAHISTE | RAHASTO |
| RAHTOURI | pseudoword | RAHTOURI | PAKOTTAVA |
| RAHVOTA | pseudoword | RAHVOTA | RAIVOTA |
| RAJUAJA | pseudoword | RAJUAJA | HOITAJA |
| RIETEVÄ | pseudoword | RIETEVÄ | ASETTUA |
| HAALAHKE | pseudoword | HAALAHKE | SIKÄLÄINEN |
| ROIMAAJA | pseudoword | ROIMAAJA | SEURAAJA |
| ROMPOTA | pseudoword | ROMPOTA | POMPPIA |
| ROPATSI | pseudoword | ROPATSI | HOUSUT |
| SAHISTA | pseudoword | SAHISTA | SÄÄTELY |
| SARKATA | pseudoword | SARKATA | JÄRKKYÄ |
| SEIPAKKO | pseudoword | SEIPAKKO | IRTISANOA |
| SIERÄMÄ | pseudoword | SIERÄMÄ | EPIDEMIA |
| SEIJINTO | pseudoword | SEIJINTO | ISTUNTO |
| SUIJAAJA | pseudoword | SUIJAAJA | OHJAAJA |
| SUUKAUTA | pseudoword | SUUKAUTA | POTKAISTA |
| HAAPATA | pseudoword | HAAPATA | HARJATA |
| SULMATA | pseudoword | SULMATA | HUUMATA |
| SÄLKEVÄ | pseudoword | SÄLKEVÄ | JALOSTAA |
| SÄÄRÄÄVÄ | pseudoword | SÄÄRÄÄVÄ | ALKUAIKA |
| TAAKKELI | pseudoword | TAAKKELI | IHANASTI |
| TAAROSTA | pseudoword | TAAROSTA | LAKAISTA |
| TAMOSNA | pseudoword | TAMOSNA | ILMEINEN |
| TIIHINKO | pseudoword | TIIHINKO | TUTKIMUS |
| TOUSATTA | pseudoword | TOUSATTA | TOISAALTA |
| TUHAAJA | pseudoword | TUHAAJA | TUHLATA |
| UIHKAAJA | pseudoword | UIHKAAJA | TYÖMÄÄRÄ |
| HAHTEVA | pseudoword | HAHTEVA | LÄHIPIIRI |
| URMETTU | pseudoword | URMETTU | TUNNETTU |
| NUULESTA | pseudoword | NUULESTA | MIELESTÄ |
| UURRONTA | pseudoword | UURRONTA | TOUKOKUU |

| Stimulus image | Stimulus Type | Stimulus Text | Model prediction |
|---|---|---|---|
| VASTINKI | pseudoword | VASTINKI | VASTINE |
| VATIERA | pseudoword | VATIERA | VÄLIERÄ |
| VEIKAAJA | pseudoword | VEIKAAJA | KIUSAAJA |
| VRAKATA | pseudoword | VRAKATA | VIRKATA |
| VÄRVÄLI | pseudoword | VÄRVÄLI | AIKAVÄLI |
| VÄTIERÄ | pseudoword | VÄTIERÄ | VÄLIERÄ |
| ÄHKÄÄJÄ | pseudoword | ÄHKÄÄJÄ | VARAAJA |
| AHDISTUS | word | AHDISTUS | AHDISTUS |
| ETEINEN | word | ETEINEN | ETEINEN |
| ÄLYKKYYS | word | ÄLYKKYYS | ÄLYKKYYS |
| ANNOSTUS | word | ANNOSTUS | ANNOSTUS |
| HUOMINEN | word | HUOMINEN | HUOMINEN |
| KORISTUS | word | KORISTUS | KORISTUS |
| KORTISTO | word | KORTISTO | KORTISTO |
| KUULUTUS | word | KUULUTUS | KUULUTUS |
| LAASTARI | word | LAASTARI | LAASTARI |
| MASENNUS | word | MASENNUS | MASENNUS |
| HUVITUS | word | HUVITUS | HUVITUS |
| LIIKUNTA | word | LIIKUNTA | LIIKUNTA |
| HARKINTA | word | HARKINTA | HARKINTA |
| HYLLYSTÖ | word | HYLLYSTÖ | HYLLYSTÖ |
| RUOHIKKO | word | RUOHIKKO | RUOHIKKO |
| VALTIAS | word | VALTIAS | VALTIAS |
| VASTAUS | word | VASTAUS | VASTAUS |
| HERTTUA | word | HERTTUA | HERTTUA |
| GORILLA | word | GORILLA | GORILLA |
| KÖYNNÖS | word | KÖYNNÖS | KÖYNNÖS |
| SUTKAUS | word | SUTKAUS | SUTKAUS |
| MAALARI | word | MAALARI | MAALARI |
| HAVAINTO | word | HAVAINTO | HAVAINTO |
| HEDELMÄ | word | HEDELMÄ | HEDELMÄ |
| HEIKKOUS | word | HEIKKOUS | HEIKKOUS |
| HELPOTUS | word | HELPOTUS | HELPOTUS |
| HEVONEN | word | HEVONEN | HEVONEN |
| HOITAJA | word | HOITAJA | HOITAJA |
| HOTELLI | word | HOTELLI | HOTELLI |
| IHMISYYS | word | IHMISYYS | IHMISYYS |
| AJATTELU | word | AJATTELU | AJATTELU |
| ILLUUSIO | word | ILLUUSIO | ILLUUSIO |
| INNOSTUS | word | INNOSTUS | INNOSTUS |
| JULISTE | word | JULISTE | JULISTE |
| JÄNNITYS | word | JÄNNITYS | LÄMMITYS |
| KAAPELI | word | KAAPELI | KAAPELI |
| KAHVILA | word | KAHVILA | KAHVILA |
| KALUSTE | word | KALUSTE | KALUSTE |
| KANERVA | word | KANERVA | KANERVA |
| KANSLIA | word | KANSLIA | KANSLIA |
| KARTANO | word | KARTANO | KARTANO |
| ANTIIKKI | word | ANTIIKKI | ANTIIKKI |
| KEISARI | word | KEISARI | KEISARI |
| KEITTIÖ | word | KEITTIÖ | KEITTIÖ |

| Stimulus image | Stimulus Type | Stimulus Text | Model prediction |
|---|---|---|---|
| KELLARI | word | KELLARI | KELLARI |
| KIRJAIN | word | KIRJAIN | KIRJAIN |
| KORAANI | word | KORAANI | KORAANIN |
| KULISSI | word | KULISSI | KULISSI |
| KÄRSIMYS | word | KÄRSIMYS | KÄRSIMYS |
| LAITURI | word | LAITURI | LAITURI |
| LEHTORI | word | LEHTORI | LEHTORI |
| LEIJONA | word | LEIJONA | LEIJONA |
| ARVOITUS | word | ARVOITUS | ARVOITUS |
| LOGIIKKA | word | LOGIIKKA | LOGIIKKA |
| LUOMINEN | word | LUOMINEN | TUOMINEN |
| LÄHETTI | word | LÄHETTI | LÄHETTI |
| MAALAUS | word | MAALAUS | MAALAUS |
| MEKLARI | word | MEKLARI | MEKLARI |
| MENESTYS | word | MENESTYS | MENESTYS |
| METALLI | word | METALLI | METALLI |
| MITTARI | word | MITTARI | MITTARI |
| MYYMÄLÄ | word | MYYMÄLÄ | MYYMÄLÄ |
| NAUTINTO | word | NAUTINTO | NAUTINTO |
| ARVOSTUS | word | ARVOSTUS | ARVOSTUS |
| NAVETTA | word | NAVETTA | NAVETTA |
| NOVELLI | word | NOVELLI | NOVELLI |
| OIKEUTUS | word | OIKEUTUS | OIKEUTUS |
| OIVALLUS | word | OIVALLUS | OIVALLUS |
| OLEMINEN | word | OLEMINEN | OSAAMINEN |
| PAKETTI | word | PAKETTI | PAKETTI |
| PALATSI | word | PALATSI | PALATSI |
| PANIIKKI | word | PANIIKKI | PANIIKKI |
| PANTTERI | word | PANTTERI | PANTTERI |
| PELAAJA | word | PELAAJA | PELAAJA |
| AURINKO | word | AURINKO | AURINKO |
| PERSOONA | word | PERSOONA | PERSOONA |
| PETTYMYS | word | PETTYMYS | PETTYMYS |
| POHDINTA | word | POHDINTA | POHDINTA |
| PRINSSI | word | PRINSSI | PRINSSI |
| PYRKIMYS | word | PYRKIMYS | PYRKIMYS |
| REALISMI | word | REALISMI | REALISMI |
| REHTORI | word | REHTORI | REHTORI |
| ROBOTTI | word | ROBOTTI | ROBOTTI |
| SAAVUTUS | word | SAAVUTUS | SAAVUTUS |
| SIVISTYS | word | SIVISTYS | SIVISTYS |
| AVOIMUUS | word | AVOIMUUS | AVOIMUUS |
| STADION | word | STADION | STADION |
| SUUNTAUS | word | SUUNTAUS | SUUNTAUS |
| SYMPATIA | word | SYMPATIA | SYMPATIA |
| TAIPUMUS | word | TAIPUMUS | TAIPUMUS |
| TARJONTA | word | TARJONTA | TARJONTA |
| TARKKUUS | word | TARKKUUS | TARKKUUS |
| TELAKKA | word | TELAKKA | TELAKKA |
| TOIVOMUS | word | TOIVOMUS | TOIVOMUS |
| TULKINTA | word | TULKINTA | TULKINTA |
| TUNNELI | word | TUNNELI | TUNNELI |
| EKONOMI | word | EKONOMI | EKONOMI |
| TUNNELMA | word | TUNNELMA | TUNNELMA |
| TUNTEMUS | word | TUNTEMUS | TUNTEMUS |
| TUNTURI | word | TUNTURI | TUNTURI |
| TUOMARI | word | TUOMARI | TUOMARI |
| TUPAKKA | word | TUPAKKA | TUPAKKA |
| TYYDYTYS | word | TYYDYTYS | TYYDYTYS |
| UPSEERI | word | UPSEERI | UPSEERI |
| UUDISTUS | word | UUDISTUS | UUDISTUS |
| VAATIMUS | word | VAATIMUS | VAATIMUS |
| VAKAUMUS | word | VAKAUMUS | VAKAUMUS |
| ELÄMINEN | word | ELÄMINEN | VALLANKUMOUS |
| VAKOOJA | word | VAKOOJA | VAKOOJA |
| VALVONTA | word | VALVONTA | VALVONTA |
| VARASTO | word | VARASTO | VARASTO |
| VARTIJA | word | VARTIJA | VARTIJA |
| VEISTOS | word | VEISTOS | VEISTOS |
| VERTAILU | word | VERTAILU | VERTAILU |
| VIRASTO | word | VIRASTO | VIRASTO |
| VOIMALA | word | VOIMALA | VOIMALA |
| YMMÄRRYS | word | YMMÄRRYS | YMMÄRRYS |
| YSTÄVYYS | word | YSTÄVYYS | YSTÄVYYS |
| [illegible] | symbols | | ALKUVAIHE |
| [illegible] | symbols | | POISSULKEA |
| [illegible] | symbols | | TILAISUUS |
| [illegible] | symbols | | LIHASKUNTO |
| [illegible] | symbols | | VERISUONI |
| [illegible] | symbols | | PURJEHDUS |
| [illegible] | symbols | | AIKATAULU |
| [illegible] | symbols | | APURAHA |
| [illegible] | symbols | | TULEHDUS |
| [illegible] | symbols | | PERIAATE |
| [illegible] | symbols | | TOUKOKUU |
| [illegible] | symbols | | JÄRJESTÄJÄ |
| [illegible] | symbols | | PURJEHDUS |
| [illegible] | symbols | | JÄLKIRUOKA |
| [illegible] | symbols | | PUUTARHA |
| [illegible] | symbols | | VALKOSIPULI |
| [illegible] | symbols | | POISSULKEA |
| [illegible] | symbols | | EPÄSUORA |
| [illegible] | symbols | | JULKISUUS |
| [illegible] | symbols | | ALUSTAVA |
| [illegible] | symbols | | JÄÄKAUSI |
| [illegible] | symbols | | PROJEKTI |
| [illegible] | symbols | | LUPAAVA |
| [illegible] | symbols | | LIPSAHTAA |
| [illegible] | symbols | | VELJEKSET |
| [illegible] | symbols | | VUOSIAIKA |
| [illegible] | symbols | | IDEOLOGIA |
| [illegible] | symbols | | KUVAPUOLI |

| Stimulus image | Stimulus Type | Stimulus Text | Model prediction |
|---|---|---|---|
| [illegible] | symbols | | EPÄSUORA |
| [illegible] | symbols | | PURJEHDUS |
| [illegible] | symbols | | LAUSUNTO |
| [illegible] | symbols | | LAUSAHDUS |
| [illegible] | symbols | | HALVEKSIA |
| [illegible] | symbols | | PURJEHDUS |
| [illegible] | symbols | | TAKAJALKA |
| [illegible] | symbols | | PURJEHDUS |
| [illegible] | symbols | | FILOSOFIA |
| [illegible] | symbols | | ITSEASIASSA |
| [illegible] | symbols | | LEPOPÄIVÄ |
| [illegible] | symbols | | LOOGISESTI |
| [illegible] | symbols | | PURJEHDUS |
| [illegible] | symbols | | PURJEHDUS |
| [illegible] | symbols | | RIPSIVÄRI |
| [illegible] | symbols | | AIHEALUE |
| [illegible] | symbols | | POSEERATA |
| [illegible] | symbols | | RATKAISEVASTI |
| [illegible] | symbols | | KERTAKAIKKIAAN |
| [illegible] | symbols | | PUUTARHA |
| [illegible] | symbols | | TARPEEKSI |
| [illegible] | symbols | | EPÄTOIVO |
| [illegible] | symbols | | KAUKOLÄMPÖ |
| [illegible] | symbols | | LOOGISESTI |
| [illegible] | symbols | | VERTAILLA |
| [illegible] | symbols | | IDEOLOGIA |
| [illegible] | symbols | | HAKUSANA |
| [illegible] | symbols | | TALVISOTA |
| [illegible] | symbols | | JOULUKUU |
| [illegible] | symbols | | LOPPUOSA |
| [illegible] | symbols | | LOOGISESTI |
| [illegible] | symbols | | UUTISOIDA |
| [illegible] | symbols | | PELIKIRJA |
| [illegible] | symbols | | POISSULKEA |
| [illegible] | symbols | | LIIKEVAIHTO |
| [illegible] | symbols | | HAALISTUA |
| [illegible] | symbols | | PAHEKSUA |
| [illegible] | symbols | | WEBSIVU |
| [illegible] | symbols | | ILOISESTI |
| [illegible] | symbols | | JÄÄKIEKKO |
| [illegible] | symbols | | VASTAUS |
| [illegible] | symbols | | BIOLOGIA |
| [illegible] | symbols | | ABSURDI |
| [illegible] | symbols | | PUOLISO |
| [illegible] | symbols | | ULKOASU |
| [illegible] | symbols | | ALKUAIKA |
| [illegible] | symbols | | POISSAOLO |
| [illegible] | symbols | | SUOJAUS |
| [illegible] | symbols | | TULIPALO |
| [illegible] | symbols | | YLÄKERTA |
| [illegible] | symbols | | KESÄPÄIVÄ |

| Stimulus image | Stimulus Type | Stimulus Text | Model prediction |
|---|---|---|---|
| [illegible] | symbols | | OLKAPÄÄ |
| [illegible] | symbols | | BROILERI |
| [illegible] | symbols | | ABSURDI |
| [illegible] | symbols | | LUOSTARI |
| [illegible] | symbols | | IDEOIDA |
| [illegible] | symbols | | TARJONTA |
| [illegible] | symbols | | PARISKUNTA |
| [illegible] | symbols | | HUOLTAJA |
| [illegible] | symbols | | PÄÄSYKOE |
| [illegible] | symbols | | IHASTUS |
| [illegible] | symbols | | PALKKATYÖ |
| [illegible] | symbols | | PURISTAA |
| [illegible] | symbols | | TOISTUA |
| [illegible] | symbols | | EPÄSUORA |
| [illegible] | symbols | | HULLUUS |
| [illegible] | symbols | | TAIDOKAS |
| [illegible] | symbols | | ULKOASU |
| [illegible] | symbols | | ULKOPUOLI |
| [illegible] | symbols | | LEPOPÄIVÄ |
| [illegible] | symbols | | BIOLOGIA |
| [illegible] | symbols | | LUPAAVA |
| [illegible] | symbols | | OIKEASSA |
| [illegible] | symbols | | LAUSUNTA |
| [illegible] | symbols | | ULKOASU |
| [illegible] | symbols | | JULKAISU |
| [illegible] | symbols | | KILPAILU |
| [illegible] | symbols | | PUSERO |
| [illegible] | symbols | | ALAMÄKI |
| [illegible] | symbols | | JULKISUUS |
| [illegible] | symbols | | BIOLOGIA |
| [illegible] | symbols | | ABSURDI |
| [illegible] | symbols | | LEHDISTÖ |
| [illegible] | symbols | | SATSATA |
| [illegible] | symbols | | ILLUUSIO |
| [illegible] | symbols | | TEOLOGIA |
| [illegible] | symbols | | LASTATA |
| [illegible] | symbols | | LÄPÄISTÄ |
| [illegible] | symbols | | ERÄPÄIVÄ |
| [illegible] | symbols | | ITÄPUOLI |
